# Dynamic integration of enteric neural stem cells in ex vivo organotypic colon cultures

**DOI:** 10.1101/2020.06.12.147652

**Authors:** Georgina Navoly, Conor J. McCann

## Abstract

Enteric neural stem cells (ENSC) have been identified as a possible treatment for enteric neuropathies. After *in vivo* transplantation, ENSC and their derivatives have been shown to engraft within colonic tissue, migrate and populate endogenous ganglia, and functionally integrate with the enteric nervous system. However, the mechanisms underlying the integration of donor ENSC, in recipient tissues, remains unclear. Here, using a modified *ex vivo* organotypic culture system we show that donor ENSC-derived cells integrate across the colonic wall in a dynamic fashion, across a three-week period. We further show that donor cells display two integrative patterns; longitudinal migration and medial invasion which allow donor cells to populate colonic tissue. Moreover, we demonstrate that significant remodelling of the intestinal ECM, and musculature, occurs upon transplantation to facilitate donor cell integration. Thus, our results provide critical evidence on the timescale, and mechanisms, which regulate donor ENSC integration within recipient gut tissue.

## Introduction

Loss of neurons within the enteric nervous system (ENS) can impact nearly every region of the gastrointestinal tract, resulting in a wide variety of disorders commonly termed enteric neuropathies.^1–6^ Such enteric neuropathies arise developmentally via disrupted development of the enteric nervous system (ENS), or postnatally via specific neuronal loss or the disturbance of neuronal signalling. Current interventions for the treatment of these diseases are mainly focused on symptom management and are limited to chronic pharmacological treatment, or surgical resection of the affected regions in the most severe cases.^7, 2^ Unfortunately, in a significant proportion of patients such interventions result in significant morbidity, and poor prognosis,^8,9^ with patients often requiring further surgical management through early childhood and adolescence.^10–13^ Given the failure of currently available surgical techniques and drug regimens to provide adequate treatment for such conditions, alternative therapeutic approaches are required.

Recently, significant research efforts have been employed to investigate the potential of autologous enteric neural stem cells (ENSC) as a possible treatment option to replace lost or damaged neurons in a range of mouse models. Early proof-of-principle studies have established the potential for *in vivo* transplantation of ENSC-derived neurons in wild-type^14–16^ and dysmotile transgenic tissues.^17, 18^ Importantly, these studies have shown the successful long-term engraftment of ENSC and their derivatives within the colonic *muscularis.* Interestingly, donor-derived cells have been observed to engraft and extend processes at the site of transplantation, forming anastomosing networks of donor cells within host tissues, which appear to functionally integrate with the endogenous ENS. Moreover, donor-derived neurons have been observed at considerable distances from the presumptive site of transplantation, and appear able to migrate through the *muscularis* to reside within endogenous ganglia structures at the level of the myenteric plexus.

However, how transplanted cells integrate into host tissues is currently unclear. Here we show, using an *ex vivo* organotypic culture system, that integration of ENSC-derived cells within myenteric ganglia occurs across a three-week timeframe. We further demonstrate that such integration requires the dynamic remodelling of the extracellular matrix (ECM), and tissue architecture, as donor cells migrate across the gut wall. Thus, we propose that these processes are rate limiting factors in the successful functional integration of ENSC-derived neurons within transplanted tissues, and are therefore fundamental considerations in the successful implementation of ENSC-based therapeutic approaches to treat enteric neuropathies.

## Results

### Characterisation of *ex vivo* organotypic cultured colon

To assess the integration of ENSC, within intestinal segments, we developed an *ex vivo* organotypic culture model in which colonic tissues could be maintained *in situ* with limited contractile forces (Fig. 1A-D), which typically lead to tissue damage when pinned. Macroscopic images of tissue segments demonstrate well-preserved, undamaged, tissue structures at day 7, 14 and 21 (Fig. 1E-F). Remarkably, up to 21 days in culture such preparations did not display evidence of culture contamination, visible structural abnormalities or any signs of detrimental tissue-scaffold interaction.

**Figure 1.**
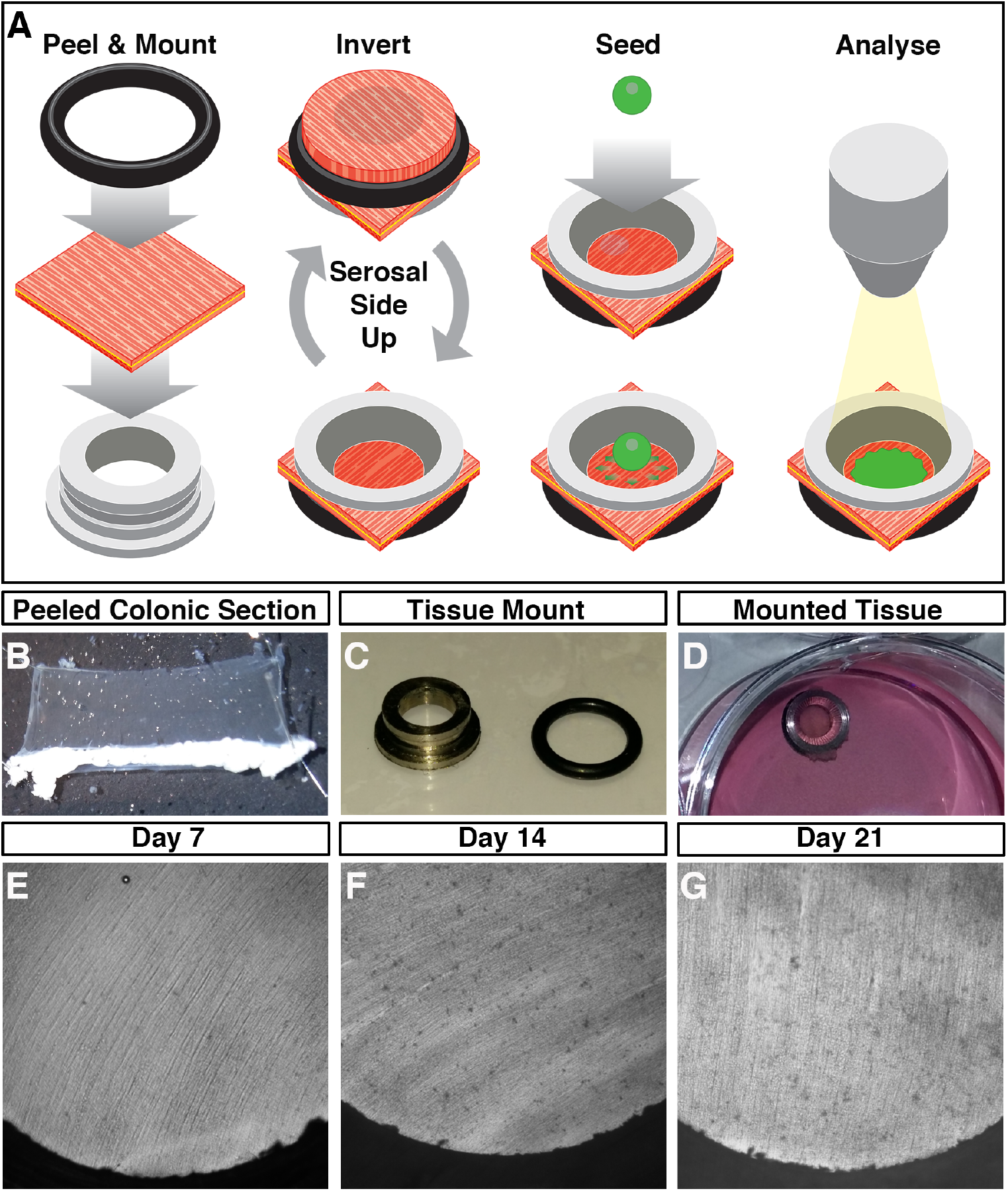
*Ex vivo* organotypic culture of colonic tissue. **(A)** Schematic representation of *ex vivo* gut culture methodology and the strategy used to perform ENSC transplantation. **(B)** Representative image of colonic *tunica muscularis* after fine dissection of the mucosal layer. **(C)** Representative image of tissue mounting apparatus. **(D)** Representative image of mounted colonic tissue segment in organotypic culture. **(E-G)** Representative images of cultured colonic tissues at Day 7 (*E*), Day 14 (*F*), and Day 21 (*G*) demonstrating the absence of contamination, and lack of gross tissue damage, within long-term organotypic cultures.

To assess the impact of organotypic culture on the development and maturation of the ENS, immunohistochemistry was performed. Within freshly dispersed (uncultured) colonic tissues, robust neuron-specific class III beta-tubulin positive (TuJ1^+^) neuronal networks could be observed (Figure 2A). TuJ1^+^ neuronal networks where observed, organised in a characteristic mesh-like structure, at the level of the myenteric plexus, in close apposition to glial fibrillary acidic protein positive (GFAP^+^) glial cells (Figure 2A, arrows). Additional TuJ1^+^ intramuscular neurons where observed extending bipolar neuronal fibres within the *tunica muscularis* (Figure 2a, arrowheads). Notably, organotypically cultured tissues demonstrated similar TuJ1^+^ and GFAP^+^ expression at day 7 (D7; Figure 2B), day 14 (D14; Figure 2C), and day 21 (D21; Figure 2D), in culture. Again, Tuj1^+^ neuronal populations could be observed both at the level of the myenteric plexus (arrows) and intramuscularly (arrowheads).

**Figure 2.**
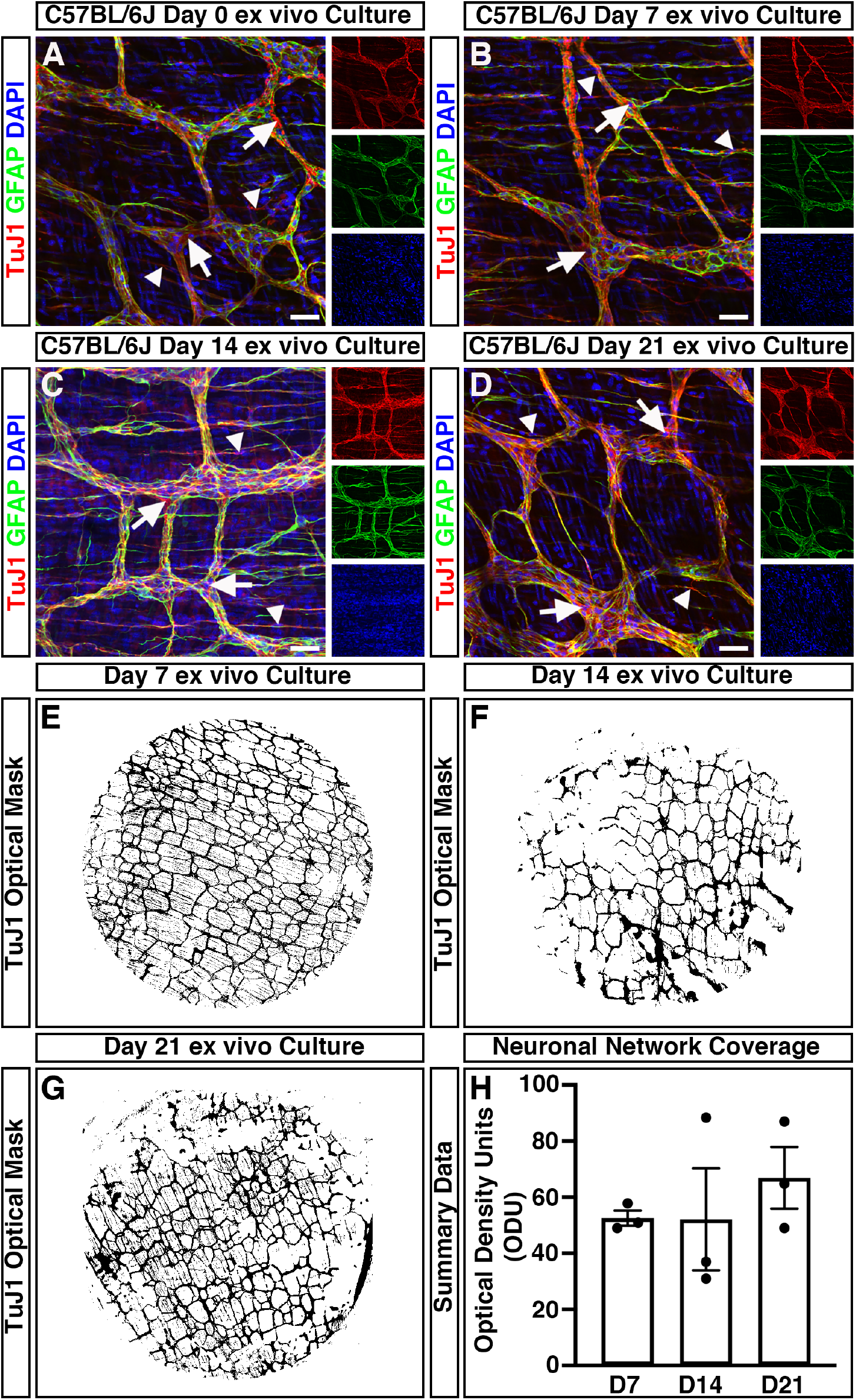
Mouse colonic segments cultured *ex vivo* maintain expression of ENS markers. **(A-D)** Representative confocal z-stack images demonstrating the presence of the pan neuronal marker; TuJ1 (*red*), glia marker; GFAP (*green*) and DAPI (*blue*) in fresh (*A*), Day 7 (*B*), Day 14 (*C*), and Day 21 (*D*) *ex vivo* cultured C57BL/6J colonic tissue. TuJ1^+^ expression was readily observed across organotypic culture and was found to label the ENS at the level of the myenteric plexus (arrows) and intramuscular nerves (arrowheads). **(E-G)** Representative optical density masks of montaged TuJ1^+^ expression within cultured colonic segments at Day 7 (*E*), Day 14 (*F*), and Day 21 (*G*). **(H)** Summary data of neuronal network coverage, as determined by the optical density of TuJ1^+^ expression, across organotypic culture (n=3 for each timepoint). Error bars represent mean ± s.e.m. Scale bars represent 50μm.

To quantitatively assess the condition of the endogenous enteric neural network, and to define its structure over time in organotypic culture, the optical density of TuJ1^+^ positive pixels within wholemount montages of cultured colonic specimens was assessed. The mean values of the optical density of TuJ1^+^ neuronal networks between D7 (52.5 ± 2.7 optical density units (ODU); n=3, Fig 2E, H), D14 (52.1 ± 18.2 ODU; n=3; Fig 2F, H) and D21 (66.9 ± 11.0 ODU; n=3; Fig 2G, H) cultures where comparable (W_2.0, 2.9_=0.66, P=0.582) as determined by Welch’s one way analysis of variance (ANOVA). These data suggest the preservation of neuronal networks within *ex vivo* organotypically cultured specimens up to 21 days in culture.

### Donor ENSC migrate extensively within organotypic cultures

ENSC were isolated from donor *Wnt1^cre/+^;R26R^YFP/YFP^* mice (P5-P7), in which neural crest cells and their derivatives express endogenous yellow fluorescent protein (YFP). This endogenous YFP expression allowed isolation, by fluorescence activated cell sorting (FACS), and fate-labelling of donor ENSC (Fig. 3A&B). Typically, enzymatic digestion of the *tunica muscularis,* from the entire small intestine and colon, and subsequent FACS lead to the enrichment of 3.2×10^5^ ± 5×10^4^ YFP^+^ cells per sorting experiment (Fig. 3B; n=5). Upon culture, individually sorted YFP^+^ cells appeared as sparsely distributed spherical shaped cells (Fig. 3C, arrows). By D3, selected YFP^+^ cells developed a typical multipolar neural crest cell appearance and began to form cellular connections (Fig. 3D, arrows). At D7, cell clusters (Fig. 3E, arrows) and elongated interconnected “chains” of cells could be observed forming a network-like structure (Fig. 3E, arrowheads). Further culture to D9 resulted in the expansion of YFP^+^ ENSC cell clusters to form extensive networks which displayed interconnecting filaments (Fig. 3F, arrowheads). Subsequently, selected YFP^+^ ENSC-derived cells were found to form multiple characteristic three dimensional “neurospheres” by approximately 2 weeks in culture (Figs. 3G&H, arrows).

**Figure 3.**
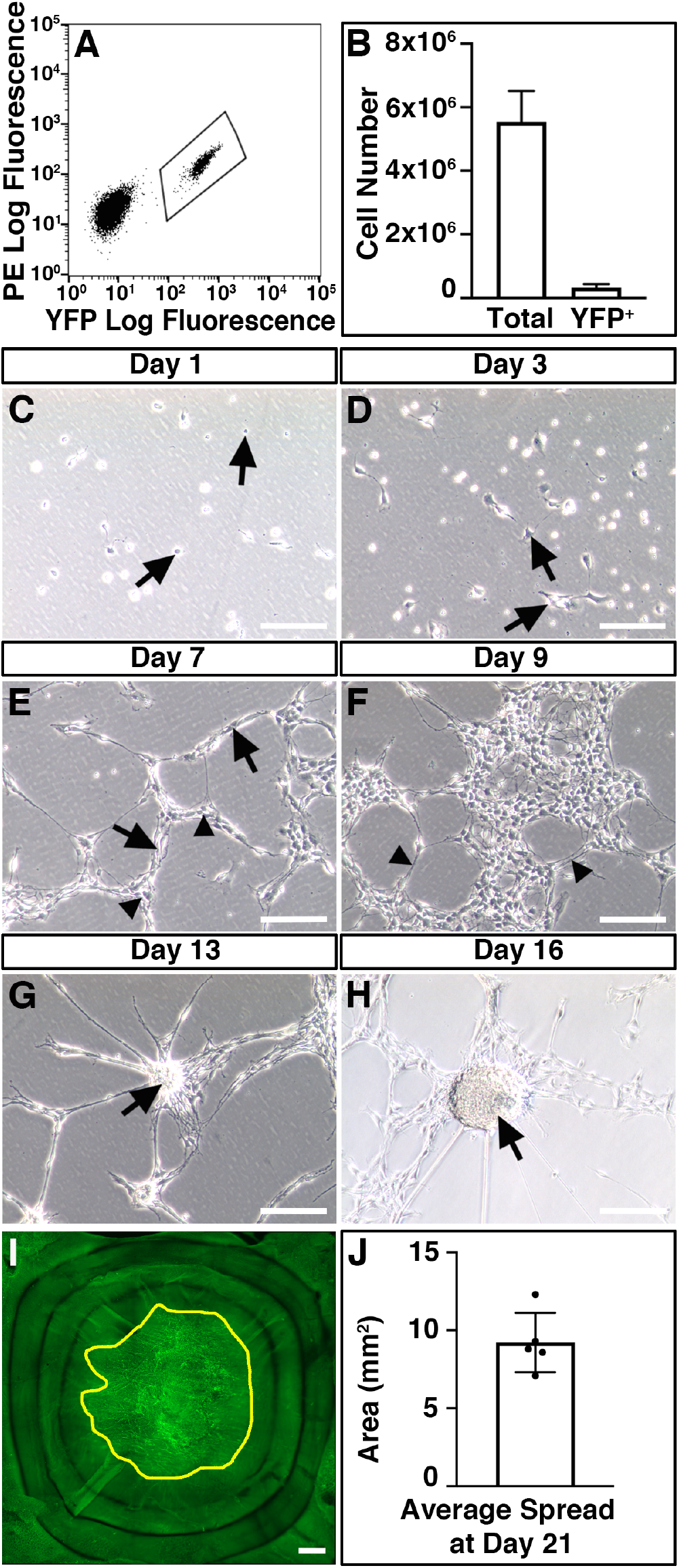
Isolation and expansion of donor ENSC for *ex vivo* transplantation. **(A)** Representative FACS profile demonstrating the gating scheme for isolation of YFP^+^ cells (bounded area) from *Wnt1^cre/+^;R26R^YFP/YFP^* intestine. **(B)** Summary data showing YFP^+^ cells populations isolated via FACS compared to total cell number. **(C-H)** Representative images showing the development of FACS-sorted YFP^+^ cells in culture. At Day 1 (*C*), the majority of sorted YFP^+^ cells displayed a spherical appearance (arrows). By Day 3 (*D)*, YFP^+^ cells developed a typical multipolar neural crest cell appearance (arrows). At Day 7 in culture (*E*), sorted cells appear as multipolar cell clusters (arrows) which are connected in a network-like structure (arrowheads). At Day 9 (*F*), YFP^+^ cell clusters appeared to form extensive networks which displayed interconnecting filaments (arrowheads). Further culture to Day 13 (*G*) and Day 16 (*H*) led to the development of rounded “neurosphere” structures (arrows). (*I*) Representative montage fluorescent image showing the development of YFP^+^ donor cells within recipient C57BL/6J colonic tissue 21-days after *ex vivo* transplantation. Yellow line bounds YFP^+^ donor cell coverage. (*J*) Summary data showing average donor cell spread within recipient tissues at Day 21. Error bars represent mean ± s.e.m (*B, J)*. Scale bars represent 250μm (*C-H*), 50 μm (*I*).

In order to assess integration of ENSC-derived cells, in intestinal specimens, we sought to establish the ability of cultured neurospheres to colonise and integrate, within organotypically cultured C57BL/6J colonic specimens *ex vivo*.

We transplanted a single, individual YFP^+^ neurosphere (approximately 2×10^4^ cells) to the serosal surface of organotypically cultured C57BL/6J colonic specimens (Fig. 3 Supplement A). At day one (D1) in culture, neurospheres appeared to be attached to recipient C57BL/6J colonic sections and multiple YFP^+^ cells were found to have dissociated from the presumptive site of transplantation, which were observed migrating as individual cells at the periphery of the donor neurosphere (Fig. 3 Supplement B). After three weeks (D21) in culture, neurosphere-derived donor YFP^+^ cells were found to have migrated extensively to cover an average area of 9.22mm² ± 0.85mm² (approximately 47% ± 4%) within recipient colonic tissues (Figs. 3I&J).

### Temporal integration of ENSC-derived cells within organotypic cultures

To assess integration of YFP^+^ donor cells, within the *tunica muscularis* of recipient colonic segments, immunohistochemistry was performed using the pan-neuronal marker TuJ1 and green fluorescent protein (GFP) antibody.

At D7, YFP^+^ cells were observed on the serosal aspect of transplanted colonic segments and appeared to migrate in multiple directions (Fig. 4A). At this stage, no penetration or integration of donor YFP^+^ cells, within the *tunica muscularis*, was observed. Donor cells were observed on the serosal aspect only (Fig. 4B; arrow), with no observable migration toward the endogenous TuJ1^+^ ENS at the level of the myenteric plexus (Fig. 4B; arrowhead)

**Figure 4.**
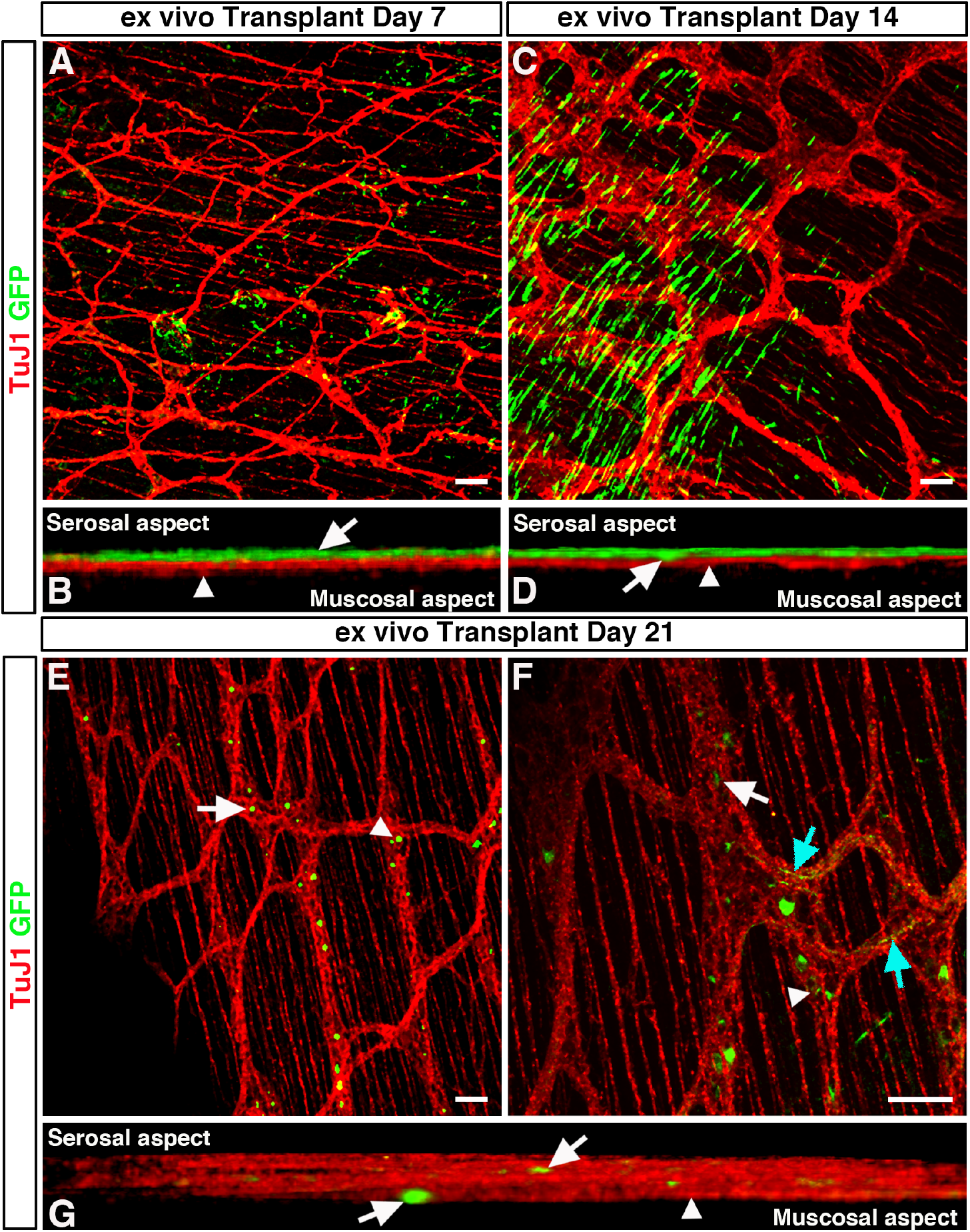
Temporal integration of transplanted ENSC within *ex vivo* organotypically cultured C57BL/6J colon. **(A)** Representative confocal z-stack image of GFP^+^ (*green*) ENSC and the endogenous TuJ1^+^ (*red*) ENS 7-days after *ex vivo* transplantation. **(B)** Orthogonal view of 3D rendered confocal stack showing donor GFP^+^ cells at the serosal aspect (*arrow)* and TuJ1^+^ myenteric plexus (arrowhead) 7-days after *ex vivo* transplantation. **(C)** Representative confocal z-stack image of GFP^+^ (*green*) ENSC and the endogenous TuJ1^+^ (*red*) ENS 7-days after *ex vivo* transplantation. **(D)** Orthogonal view of 3D rendered confocal stack, 14-days after *ex vivo* transplantation, showing donor GFP^+^ cell penetration (*arrow*) from the serosal aspect towards the myenteric plexus (arrowhead). **(E&F)** Representative low-(*E*), and high-power (*F*) confocal z-stacked images of GFP^+^ (*green*) donor cells within the recipient TuJ1^+^ (*red*) myenteric plexus, 21-days after *ex vivo* transplantation. Donor cells were observed as both individual cells (arrowheads), and donor cell clusters (arrows), which appeared to extend GFP^+^ fibres which traced the endogenous neural network (cyan arrows). **(G)** Orthogonal view of 3D rendered confocal stack showing donor GFP^+^ cells (*arrows)* within the endogenous TuJ1^+^ myenteric plexus (arrowhead) 21-days after *ex vivo* transplantation. Scale bars represent 50μm.

At post-transplantation D14, transplanted YFP^+^ neurospheres appeared to have dissociated into single cells (Fig. 4C) with numerous, spindle shaped, YFP^+^ cells observed. The majority of YFP^+^ cells at D14 were again observed at the outer serosal aspect of transplanted tissues. However, donor YFP^+^ cells were observed which displayed a limited degree of tissue penetration (Fig. 4D; arrow) towards the myenteric plexus (Fig. 4D; arrowhead).

By contrast, at D21 transplanted YFP^+^ ENSC were observed fully integrated within recipient endogenous enteric ganglia, at the level of the myenteric plexus (Fig. 4E-G). YFP^+^ cells appeared to be distributed along the “mesh-like” myenteric neural network, both as individual cells (Fig. 4E&F; arrows) and as donor cell clusters within individual ganglia structures (Fig. 4E&F; arrowheads). Moreover, YFP^+^ fibres were also observed at the level of the myenteric plexus, which appeared to trace the endogenous recipient neural network (Fig. 4F; cyan arrows). Upon three-dimensional (3D) reconstruction, donor YFP^+^ cells could clearly be observed throughout the TuJ1^+^ myenteric neural network (Fig. 4G; arrowhead), including at both the serosal and mucosal aspects of the myenteric plexus (Fig 4G; arrows).

We next aimed to quantify the maximal distance ENSC-derived cells had migrated across the gut wall using confocal z-stacks of GFP-labelled donor cells, in combination with DAPI labelling (Fig. 5A&B). Here, maximal cell integration was recorded by observing the most medially integrated donor cell body (GFP^+^/DAPI^+^), relative to the serosal surface using Z-series scans (1μm optical sections), which showed significant differences in the mean values across the three timepoints as determined by Welch’s ANOVA (W_2.0, 5.9_=5.58, P=0.04). At D7 ENSC-derived donor cells were observed at a z-axis depth of 9.5 ± 1.6 μm (Fig. 5C; n=4). Similarly, at D14 donor-derived cells were observed at a z-axis depth of 9.3 ± 1.4 μm (P=0.91; n=4; Fig. 5C). However, at D21 the most medially integrated donor-derived cell body in *ex vivo* cultured transplanted colon was observed at a z-axis depth of 15 ± 1.2 μm (P=0.03 vs D7, P=0.02 vs D14; n=4; Fig. 5C). Importantly, depth-coding of all GFP^+^ donor-derived cell bodies and fibres, according to their z-axis depth, revealed integration of donor ENSC along the serosal aspect of the gut (Fig. 5D; arrowheads, Supplementary Movie 1). Moreover, from the presumptive site of transplantation (Fig. 5D; magenta arrow), donor cells appear to penetrate the gut wall at the site of neurosphere engraftment (Fig. 5D; dashed white line) and migrate along the longitudinal aspect of the gut within the *tunica muscularis* (Fig. 5D; white arrows).

**Figure 5.**
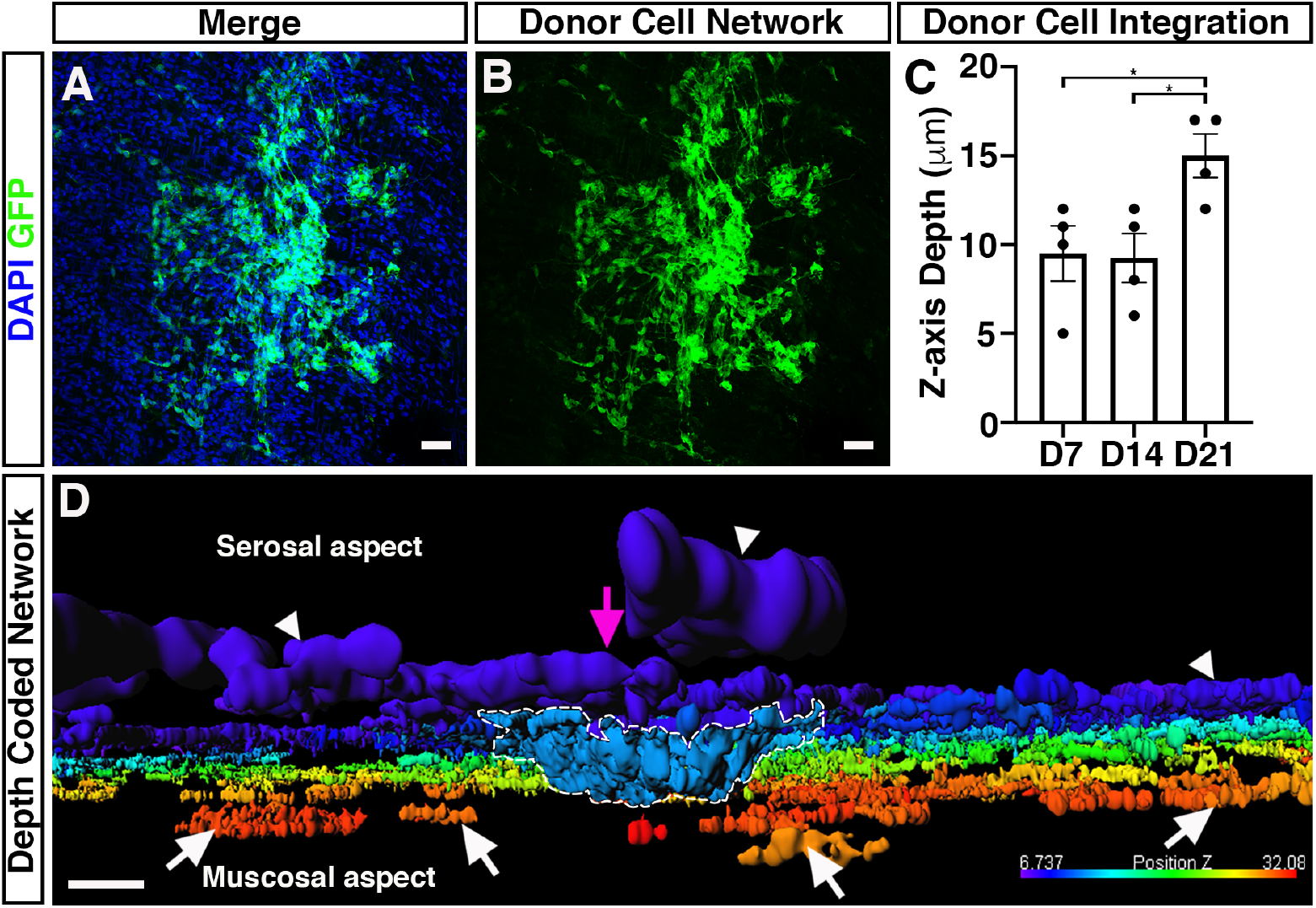
Integration of ENSC-derived donor cells across the gut wall. **(A&B)** Representative confocal z-stack image of donor ENSC-derived GFP^+^ (*green*), and DAPI (blue), within the gut wall 21-days after *ex vivo* transplantation. **(C)** Summary data showing analysis of donor cell body (GFP^+^/DAPI^+^) integration, measured as z-axis depth, across the gut wall at day-7 (D7), D14, and D21 after *ex vivo* transplantation. **(D)** Representative depth-coded image of donor cell integration at D21. Note the protrusion of donor ENSC-derived cells into the gut wall (*dashed line*) from the site of transplantation on the serosal surface (*magenta arrowhead*). Donor GFP^+^ cells and fibres were observed to have integrated longitudinally along the gut, both on the serosal surface (*arrowheads*) and within the *tunica muscularis* (*arrows)*. Scale bars represent 50μm (*A, B*), 30 μm (*D*). Error bars represent mean ± s.e.m. ^*^ P≤ 0.05, by Welch’s t-test.

Taken together, these data suggest that ENSC appear to initially migrate from the transplanted neurosphere, along the serosal aspect of the tissue, in a non-uniform manner. Subsequently, transplanted cells penetrate the *tunica muscularis* at the site of neurosphere transplantation, and migrate within the *tunica muscularis* towards the endogenous neural network at the level of the myenteric plexus, allowing integration within ganglia structures and extension of donor cell processes, within the myenteric network, by approximately three weeks.

### Molecular characterisation of donor ENSC integration after transplantation

Having demonstrated the integration of donor ENSC-derived cells, within the *tunica muscularis,* we assessed possible tissue remodelling within the gut wall; and the molecular basis of ENSC migration and integration within transplanted colonic segments.

In control (non-transplanted) tissues, the smooth muscle architecture appears to be preserved up to three weeks in organotypic culture (Fig. 6A-C). Here, immunohistochemistry for the smooth muscle marker SM22 revealed smooth muscle cells in close apposition, both in the longitudinal (Fig. 6B) and circular (Fig. 6C) orientations. Orthogonal reconstruction of confocal z-stacks also revealed intact SM22^+^ muscle layers in both the longitudinal and circular planes (Fig. 6A^i^). By contrast, in transplanted tissues, disruption of the smooth muscle architecture was observed at the site of transplantation 21-days following ENSC transplantation (Fig. 6D-G). Examination of the longitudinal muscle layers revealed complete loss of SM22^+^ expression at the site of neurosphere engraftment (Fig. 6E; white star). Similarly, examination of the circular muscle layer revealed partial reductions in SM22 expression (Fig. 6F; white star). Interestingly, this disruption in SM22 expression appeared to be limited to the site of transplantation and neurosphere engraftment. Upon orthogonal reconstruction of the engraftment site (Fig. 6D, ROI 1), GFP^+^ donor cell expression was observed within the plane of the longitudinal muscle layer (Fig. 6D^i^). While this donor cell expression appeared restricted to the serosal aspect, and presumptive longitudinal muscle layer, all SM22^+^ expression in this engraftment region was absent. However, in regions away from the site of engraftment (Fig. 6D, ROI 2) uninterrupted SM22^+^ expression was observed in both the longitudinal and circular muscle layers (Fig. 6D^ii^). Immunohistochemistry also revealed apparent alterations in the extracellular matrix (ECM) within transplanted tissues. As opposed to control non-transplanted colon, where Collagen type IV (Col IV) expression was observed in near continuity (Fig. 6H), loss of Collagen IV was observed at the boundary of the engraftment site in transplanted tissues (Fig. 6I; *magenta border*). Additionally, the presence of Collagen IV expression was observed, at the site of engraftment, within the remaining neurosphere structure (Fig. 6I; *cyan arrows*).

**Figure 6.**
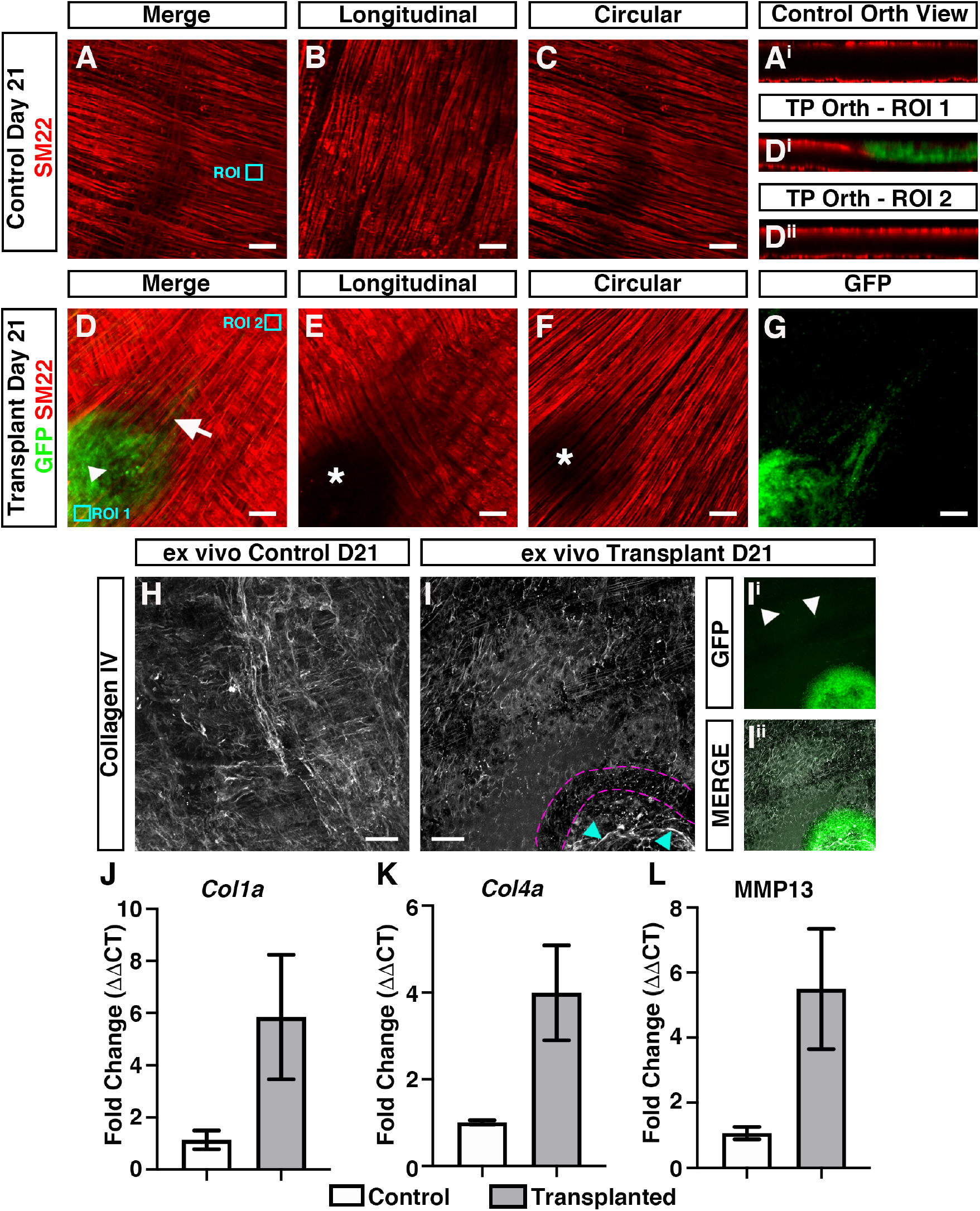
Donor cell integration induces tissue remodeling after *ex vivo* transplantation. **(A)** Representative confocal z-stack image of SM22^+^ (*red*) smooth muscle within control (non-transplanted) C57BL/6J colonic tissue at day-21. **(A^i^)** Orthogonal view taken from the region of interest (*ROI, cyan box)* in *A* showing SM22^+^ smooth muscle in two uninterrupted layers. **(B&C)** Representative images of individual z-stacks at the longitudinal (*B*) and circular (*C*) muscle layers. **(D)** Representative confocal z-stack image of donor GFP^+^ (*green*) and SM22^+^ (*red*) smooth muscle 21-days after *ex vivo* transplantation. (**D^i^&D^ii^**) Orthogonal views taken from *ROIs (cyan boxes)* in *D* showing interruption of the longitudinal smooth muscle (*D^i^*) at the site of neurosphere engraftment. At sites away from the area of neurosphere engraftment SM22^+^ expression appears to demonstrate two uninterrupted smooth muscle layers (*D^ii^*). **(E&F)** Representative images of individual z-stacks at the longitudinal (*E*) and circular (*F*) muscle layers 21-days after *ex vivo* transplantation. **(G)** Representative image of individual z-stack, at the level of the circular muscle (taken from *D*), showing migration of GFP^+^ donor derived cells from the site of neurosphere engraftment 21-days after *ex vivo* transplantation. **(H)** Representative confocal z-stack image of Collagen IV (*grey*) in control (non-transplanted) C57BL/6J colonic tissue at day-21. **(I)** Representative confocal z-stack image of Collagen IV (*grey*) 21-days following *ex vivo* transplantation. Note the presence of Collagen IV expression at the site of neurosphere transplantation (*cyan arrows*) and apparent discontinuity of Collagen IV expression surrounding the presumptive neurosphere boundary (*magenta dashed lines*). **(I^i^&I^ii^)** Representative GFP channel (*I^i^*) and merged (*I^ii^*) confocal z-stack images showing GFP^+^ donor cells at the site of transplantation and within the *tunica muscularis* (arrows) having migrated away from the engrafted neurosphere. **(J-L)** Summary data showing mRNA fold change for extracellular matrix (ECM)-related genes 21-days after *ex vivo* transplantation. ECM genes (*grey bars*) Collagen 1a (*Col1a*; *J*), Collagen 4a (*Col4a*; *K)* and Matrix metalloproteinase 13 (*MMP13; L)* were found to be expressed at increased levels compared to non-transplanted control (*white bars*) tissues. Scale bars represent 50μm. Error bars represent mean ± s.e.m.

To further examine the mechanisms involved in donor cell integration, molecular analyses were performed to examine the expression of candidate genes, known to play critical roles within the intestinal ECM, such as Collagen 1a2 (*Col1a*) and Collagen 4a1 (*Col4a*), and in remodelling of the ECM, including Matrix metalloproteinase 1 (*MMP1*), *MMP8* and *MMP13*. Of note, *MMP1* and *MMP8* could not be resolved in control postnatal C57BL/6J wild type colonic tissue, whereas *Col1a*, *Col4a* and *MMP13* were found to be robustly expressed (*data not shown*) using standard real-time polymerase chain reaction (RT-PCR) analysis. Therefore, *Col1a*, *Col4a* and *MMP13* were taken forward for quantitative RT-PCR (qRT-PCR) analysis to assess their expression in transplanted *ex vivo* tissue segments compared to control non-transplanted colon.

Interestingly, ENSC integration within transplanted tissues led to increased expression of *Col1a* (5.8 ± 2.4-fold increase; P=0.007; n=4; Fig. 6J), *Col4a* (4.0 ± 1.1-fold increase; P=0.011; n=4; Fig. 6K) and *MMP13* (5.5 ± 1.9-fold increase; P=0.008; n=4; Fig. 6L) when compared to control (non-transplanted) tissues, as determined by comparison of ΔCT values by Welch’s t-test analysis.

These data suggest that active remodelling of the ECM and smooth muscle architecture occurs in response to ENSC transplantation, which subsequently allows integration of donor ENSC within and along the gut wall.

## Discussion

Over the past decade there has been an increasing focus on the development and evaluation of stem cell-based therapies for treating enteric neuropathies. Recent studies have highlighted the potential of autologous, and pluripotent-derived ENS progenitor-based therapy, as a means of replacing neurons after *in vivo* transplantation to mouse colon.^14–17, 19–23^ However, the precise mechanisms by which donor-derived cells integrate within recipient tissue remain unclear. Therefore, studies to uncover the timeframe, and mechanisms, underlying donor cell integration are fundamental to progressing towards clinical application of stem cell therapy for gut disorders. Moreover, mechanistic studies of donor cell integration may uncover useful targets to improve the efficiency of donor cell integration, and function, within recipient tissues.

Importantly, many previous studies have relied heavily on *in vivo* surgical transplantation procedures to rodents.^14–18, 20, 24^ While this has provided crucial proof-of-principle data that donor cells *can* integrate within the gut after transplantation, technical limitations such as tissue opacity, and an inability to perform long-term *in vivo* imaging, have limited the mechanistic investigation of *how* donor cells integrate within recipient gut tissue. Alternative approaches including explant cultures have also established the feasibility of transplanting donor enteric precursors into aneural gut.^25–28^ However, these explant methods are often hindered by limitations in their reproducibility, or in the long-term maintenance of tissues on chick chorioallantoic membrane (CAM) cultures.^28–30^

To begin to overcome some of these limitations, here we have employed an *ex vivo* organotypic culture method to investigate the temporal integration of ENSC within murine gut segments. We demonstrate that this *ex vivo* organotypic culture configuration allows for long-term culture of murine colonic segments, with preservation of both tissue architecture and ENS structure, providing a robust, reproducible and readily scalable model. Significantly, this method also appears to limit tissue damage, within culture colonic segments, when compared to standard *ex vivo* organotypic culture conditions which typically require pinning of tissue. Hence, this organotypic method provides an optimal model to explore the temporal integration of donor cells within gut tissues.

Previous studies have detailed the functional integration of donor neurons, derived from postnatally harvested ENSC, in recipient wild-type colonic tissues.^14, 20^ These studies, using calcium imaging and optogenetic approaches, have detailed the functional integration of donor ENSC-derived neurons within the host neuromusculature four weeks after *in vivo* transplantation. By four weeks, engrafted cells were found to have migrated extensively within the recipient gut wall, including the integration of donor-derived cells within endogenous ganglia, and the extension of graft-derived axonal fibres which appeared to both trace the endogenous ENS structure, and project to the circular muscle layer. Moreover, engrafted donor-derived neurons were found to both functionally integrate with the endogenous ENS circuitry^14^ and provide inhibitory, and excitatory, innervation to intestinal smooth muscle.^20^ A similar timescale was also observed in the functional integration of ENSC-derived neurons within dysmotile transgenic tissues.^17^ Here, ENSC transplantation was found to restore neural responses and rescue gut transit four weeks after transplantation. Again, significant migration of ENSC-derived cells was observed both within, and along, the gut wall with donor cells appearing to “home” to the myenteric plexus. More recently, derivation of ENS progenitors from pluripotent sources, such as embryonic stem cells or induced pluripotent stem cells has been proposed as a potential “off-the-shelf” alternative to autologous ENSC.^21^ Similar to ENSC, pluripotent-derived ENS progenitors have been shown to engraft and migrate through the colonic *muscularis* to reside within endogenous ganglia structures, across a four week period, following *in vivo* transplantation.^22^ Taken together, these findings suggest that the migration of donor ENSC-derived cells into the gut neuromusculature, and the “homing” of donor neurons to the myenteric plexus region, are likely central processes in the establishment of donor-derived functional responses within the host neuromusculature. However, to date, a more definitive timeframe and mechanistic understanding of donor cell integration remains elusive.

Using our modified *ex vivo* transplantation approach, we demonstrate that migration of ENSC-derived cells away from the presumptive site of transplantation, and integration within the colonic musculature, is a dynamic process which occurs over a three-week period. Initially, ENSC-derived donor cells were observed to migrate away from the site of neurosphere engraftment, along the serosal aspect of the colon. This migration at the serosal surface did not appear to show preferential directionality and is reminiscent of the migration pattern of cultured neural crest cells.^31–33^ A secondary phase of medial migration appears to occur allowing donor cells to integrate across the gut wall. Crucially, this secondary process appears to be established over a number of weeks, with penetration of the neuromusculature, and medial migration, appearing to be prominent only after approximately 14 days.

We hypothesised that donor ENSC utilise molecular processes analogous to that of metastatic cancer cells, in order to remodel the ECM, infiltrate (i.e. intra/extravasate) and migrate (i.e. invade) within the host gut wall following transplantation. Indeed, previous studies have highlighted that the matrisome plays multiple dynamic roles in embryonic neural crest cell migration and differentiation.^33–36^ In tandem, migratory neural crest cells have been shown to modify the local ECM, suggesting that migration through the ECM is regulated on multiple levels.^37, 38^

Our initial expectation was that migration through the colonic neuromusculature, and integration in myenteric ganglia, would necessitate collagenase-based depletion of the ECM. Our finding of an upregulation in Collagenase 3 *(MMP13)* expression, within transplanted *ex vivo* cultured tissues, fits with our hypothesis as *MMP13* has been shown to be involved in the progression of colorectal cancer.^39, 40^ Importantly, MMP proteolysis of Collagen IV has previously been shown to promote cell migration, via the production of specific cleavage fragments with independent biological activity.^41, 42^ Therefore, upregulation of *MMP13* in transplanted *ex vivo* colon may provide a possible mechanism of how transplanted ENSC remodel the ECM, to allow invasion of donor cells. Of note, early emigration of neural crest cells from the neural tube has been shown to be associated with discontinuity of Collagen IV staining.^34^ Similarly, here we demonstrate that major remodelling of the musculature and Collagen IV occurs, after *ex vivo* transplantation, around the site of neurosphere engraftment which appears to be consistent with neural crest emigration after transplantation. Yet, our findings of increased Collagen (*Col1a and Col4a*) gene expression were unexpected. Previous investigations have, however, demonstrated dynamic regulation of Collagen I and IV during neural crest cell migration. In particular, Collagen I expression has been shown to be upregulated along neural crest migratory pathways, whereas Collagen IV has been shown to be increased upon neural crest cell aggregation and differentiation.^34^ Hence, these processes may account for the observations in the current study, as transplanted ENSC have been shown to migrate extensively within *ex vivo* transplanted tissue, and aggregate within endogenous ganglia structures at the level of the myenteric plexus.

While this study has revealed the timescale and a possible mechanism underlying donor cell integration within host tissues following ENSC transplantation, future studies will be required to fully elucidate the mechanisms involved. Here, we utilised early post-natal (P5-7) ENSC to investigate donor cell integration within C57BL/6J wild-type colon. A recent study has shown that enteric neural crest cells progressively lose capacity to form the ENS with age.^29^ Hence, further studies will be required to determine the integration capacity of aged donor cells. This caveat may have significant implications in the clinical translation of cellular therapies for enteric neuropathies, as donor age will need to be considered for both autologous cell transplants, and in strategies using pluripotent cell sources; given time in culture may alter integrative capacity. Importantly, the outlined *ex vivo* transplantation methodology may offer significant benefits in tackling such questions in a medium-throughput, and consistent, manner. Furthermore, we used C57BL/6J wild-type colon as recipient tissue in this proof-of-principle study. We demonstrate that donor-derived cells infiltrate and “home” to the myenteric ganglia by three weeks post-transplantation. We assume that trophic factors regulate this “homing” process, though the exact factors involved remain unclear. Moreover, recent studies have demonstrated that the colonic ECM is altered in diseased states,^43–45^ which suggests the integrative ability of ENSC-derived donor cells is likely to be altered by the recipient microenvironment in different disease states. Future studies will therefore be required to address how disease-specific parameters alter ENSC integration, which may be clinically relevant.

We conclude that this study provides critical evidence on the timescale and mechanisms which regulate ENSC integration within recipient gut tissue. Furthermore, the modified organotypic culture model utilised in the current study may be beneficial in the future assessment of donor cell transplantation in neuropathic models, or in the development of targeted approaches to influence donor cell behaviour as a possible combinatorial strategy for the treatment of enteric neuropathies.

## Materials and Methods

### Animals

Animals were obtained from The Jackson Laboratory (Bar Harbor, MN, USA). For experimental procedures, adult C57BL/6J wild-type (6-8 week old) were used to obtain recipient tissue, and early postnatal (day 5-7) *Wnt1^cre/+^;R26R^YFP/YFP^* mice (where neural crest cell derivatives express yellow fluorescent protein (YFP) were used as donors. Animals were housed and experiments were performed in accordance with the UK Animals (Scientific Procedures) Act 1986, and approved by the University College London Biological Services Ethical Review Process. Animal husbandry at UCL Biological Services was in accordance with the UK Home Office Certificate of Designation.

### Donor cell isolation and enrichment

The entire gut (small intestine and colon) was obtained from *Wnt1^cre/+^;R26R^YFP/YFP^* mice at P5-7 after cervical dislocation. Tissues were removed to sterile phosphate-buffered saline (PBS, 0.01 mol L^−1^, pH 7.2 at 4°C) for further dissection. Strips of the *tunica muscularis* were obtained from the jejunum, ileum and colon following removal of the mucosa via fine dissection.

Single intestinal cells were obtained after enzymatic dissociation of the *tunica muscularis* using a Tumor Dissociation Kit (Miltenyi Biotec, Woking, UK), and YFP^+^ cells isolated using fluorescence activated cell sorting (FACS) with a MoFloXDP cell sorter (Beckman Coulter, Wycombe, UK). YFP^+^ cells were selected using a 530/40 filter set.

### Neurosphere Culture

YFP^+^ cells were plated at a minimum seeding density of 1 × 10^5^ cells/well on fibronectin-coated (2% w/v in 0.1mol l^−1^ PBS, Sigma-Aldrich, Gillingham, UK) 6-well dishes. Plated YFP^+^ cells were maintained in “neurosphere medium” (NSM; DMEM/F12 supplemented with B27 (Gibco, Hemel Hempstead, UK), N2 (Gibco), 20ng/ml epidermal growth factor (EGF, Peprotech, London, UK), 20ng/ml fibroblast growth factor (FGF, Peprotech), and Primocin (100μg/ml; InvivoGen, Toulouse, France) antibiotic. Typically, such cultures from P5-P7 intestine formed “neurospheres” between 1-2 weeks and were maintained in culture for up to 4 weeks.

### Organotypic culture

The entire colon from adult (6-8 week old) C57BL/6J mice was removed to sterile PBS (0.01 mol L^−1^, pH 7.2 at 4°C) after cervical dislocation. Colonic tissues were pinned in a Sylgard-lined chamber and opened along the length of the colon at the mesenteric border. The mucosa was removed, via fine dissection, taking special care to avoid any damage to the underlying *tunica muscularis*. Tissue segments were then mounted (serosa side down) on sterilized 5mm diameter metal tissue mounts and fixed in place with sterilized “O-rings.” Tissues were washed thoroughly in sterile PBS (0.01 mol L^−1^, pH 7.2) supplemented with Primocin (100mg/ml; InvivoGen) and maintained in Dulbecco’s Modified Eagle Medium (DMEM, Gibco) supplemented with L-Glutamine and Primocin (100mg/ml; InvivoGen) and maintained at 37°C, 95% O_2_/5% CO_2_ for 1 hour before neurosphere transplantation.

### Ex vivo neurosphere transplantation

Colonic tissue scaffolds were inverted and placed, submucosal side down, into individual wells of an untreated 6-well plate. An individual YFP^+^ neurosphere was subsequently transplanted to the serosal surface, by mouth pipette, using a pulled glass micropipette. 15μl media (DMEM, L-Glutamine and Primocin) was added inside the scaffold wall and transplanted tissues were incubated for 2 hours at 37°C, allowing attachment of the neurosphere to the serosal surface. Subsequently, wells were supplemented with 3ml media (DMEM, L-Glutamine and Primocin) and were maintained in culture for between 1-3 weeks post-transplantation.

### Immunohistochemistry

Tissues were fixed in paraformaldehyde (4% w/v in 0.1 mol L^−1^ PBS) for 45 min at room temperature (RT). After fixation, tissues were washed thoroughly for 1h in PBS (0.01 mol L^−1^, pH 7.2 at RT). Tissues were blocked for 1h (0.1 mol L^−1^ PBS containing 1% Triton X-100, 10% sheep serum). Tissues were incubated in primary antibody (diluted in 0.1 mol L^−1^ PBS containing 1% Triton X-100, 10% sheep serum, Supplementary Table 1) for 16h at 4°C and immunoreactivity was detected using the secondary antibodies listed in Supplementary Table 2 (1:500 in 0.1 mol L^−1^ PBS, 1h at RT). Before mounting, tissues were washed thoroughly in PBS (0.1 mol L^−1^ PBS for 2h at RT). Control tissues were prepared by omitting primary or secondary antibodies. Tissues were examined using as LSM710 Meta confocal microscope (Zeiss, Munich, Germany). Confocal micrographs were digital composites of Z-series scans (0.5μm-1μm optical sections). Final images were constructed using FIJI software.^46^ Depth-coding of confocal metafiles was performed using Imaris Cell Imaging Software (Oxford Instruments, Zurich, Switzerland).

### Quantification of transplanted ENSC spread

For montage experiments of cell spread, tissues were examined using an Axioplan Observer microscope (Zeiss). Micrographs of wholemounts were digital composites of individual 20X tiled micrographs, stitched using Zen software (Zeiss). Final images were constructed using FIJI software. The area of donor cell migration within recipient colonic tissue scaffolds was quantified (Adobe Photoshop, CA, USA), with individual and mean values plotted using GraphPad Prism software (GraphPad, CA, USA).

### RT-PCR

RNA was extracted from the control and transplanted tissues using TRIzol reagent (ThermoFisher, Hemel Hempstead, UK) and treated with DNase I (Qiagen, Manchester, UK). First-strand cDNA was amplified from 100ng RNA using SuperScript^®^ VILO™ cDNA Synthesis Kit (ThermoFisher). RT-PCR was performed to initially examine the expression of candidate genes including Glyceraldehyde 3-phosphate dehydrogenase *(GAPDH), Col1a, Col4a, MMP1, MMP8 and MMP13* in control tissues. qRT-PCR was performed with a StepOnePlus Real-Time PCR System (ThermoFisher) using the Quantitect SYBR Green PCR kit (Qiagen), according to the manufacturer’s instructions. qRT-PCR was performed in triplicate, using region-specific primers designed against mouse sequences for *GAPDH, Col1a, Col4a* and *MMP13* (Supplementary Table 3). Gene expression data were expressed as a proportion of *GAPDH*, as a reference, using ΔCT and ΔΔCT calculations.

### Statistical analysis

Data are expressed as mean ± standard error of the mean. Statistical analysis was performed using GraphPad Prism software (GraphPad). The intergroup differences for neural network quantification, and maximal z-axis integration post-transplantation, at day 7, 14, & 21, were evaluated using Welch’s ANOVA and Welch’s t-tests. For transcriptomic analyses statistical comparison between control (non-transplanted) and transplanted samples was performed using ΔCT values by Welch’s t-test. Results were considered significant at P < 0.05. The ‘n values’ reported refer to the number of colonic segments, each from a separate mouse, examined at each timepoint.

### Data Availability

The data that support the findings of this study are available from the corresponding author upon reasonable request.

## Acknowledgements

The authors would like to thank Dr. Ayad Eddaoudi, (UCL Great Ormond Street Institute of Child Health Flow Cytometry Facility), Dr. Dale Moulding (UCL Great Ormond Street Institute of Child Health Imaging Facility), Chey Chapman and Ben Cairns for technical support, along with Dr. Nikhil Thapar for advice in refining the manuscript.

The authors would like to acknowledge the NIHR Great Ormond Street Hospital Biomedical Research Centre which supports all research at Great Ormond Street Hospital NHS Foundation Trust and UCL Great Ormond Street Institute of Child Health. The views expressed are those of the author(s) and not necessarily those of the NHS, the NIHR or the Department of Health.

## Author contributions

GN & CMC acquired and interpreted data. CMC obtained funding. GN & CMC contributed to study concept and design drafted and critically revised the manuscript.

---

**Figure 3 Supplement.**
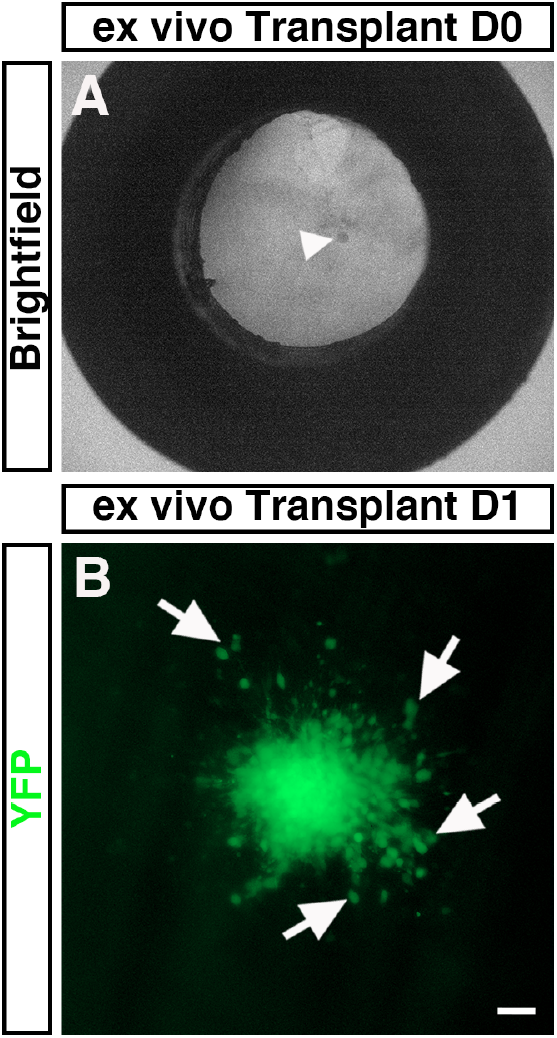
Early integration and migration of ENSC in recipient colon following *ex vivo* transplantation. **(A)** Representative low-power brightfield image, taken at transplantation, showing mounted C57BL/6J colonic tissue with transplanted neurosphere *in situ* (*arrowhead*). **(B)** Representative z-stacked image showing the migration of donor YFP^+^ ENSC-derived cells (arrows) away from the site of neurosphere integration 1-day after *ex vivo* transplantation. Scale bar represents 50μm.

**Supplementary Movie 1. Transplanted ENSC integrate extensively across the colonic wall following** *ex vivo* **transplantation.** Video (*15 fps*) of depth-coded confocal micrograph demonstrating integration of ENSC-derived GFP^+^ donor cells across the gut wall, in C57BL/6J colonic tissue, 21-days after *ex* vivo transplantation. Initially, GFP^+^ donor cells can be seen to have migrated extensively, in both the longitudinal and circumferential directions, from the engraftment site. Depth-coding of this GFP^+^ network shows the integration of ENSC-derived GFP^+^ cells at variable depths within the *tunica muscularis*.

**Supplementary Table 1.**
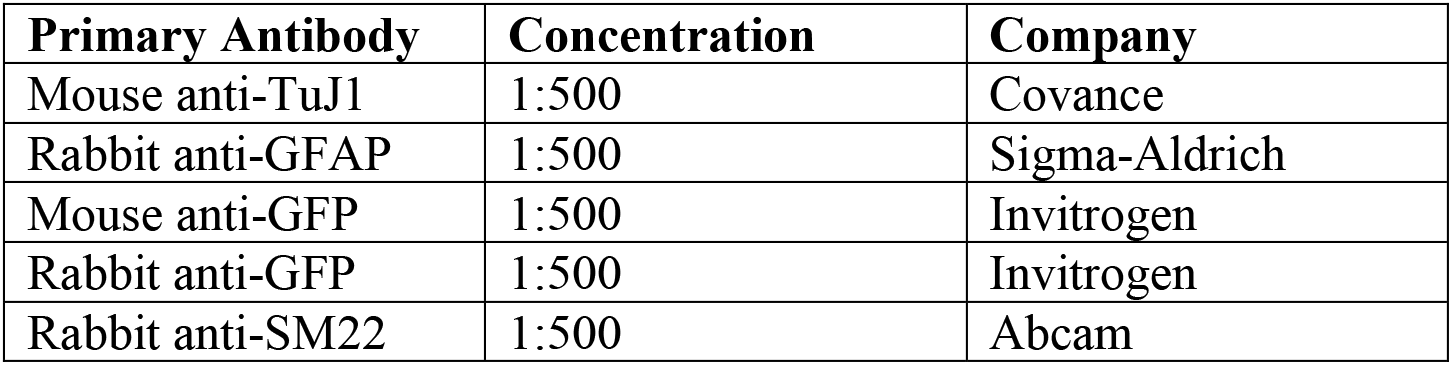
Primary Antibodies used for Immunohistochemistry

| Primary Antibody | Concentration | Company |
| --- | --- | --- |
| Mouse anti-TuJ1 | 1:500 | Covance |
| Rabbit anti-GFAP | 1:500 | Sigma-Aldrich |
| Mouse anti-GFP | 1:500 | Invitrogen |
| Rabbit anti-GFP | 1:500 | Invitrogen |
| Rabbit anti-SM22 | 1:500 | Abcam |

**Supplementary Table 2.** Secondary Antibodies used for Immunohistochemistry

| Secondary Antibody | Alexa Fluor | Concentration | Company |
| --- | --- | --- | --- |
| Goat anti-mouse | 488 | 1:500 | Invitrogen |
| Goat anti-mouse | 568 | 1:500 | Invitrogen |
| Goat anti-rabbit | 488 | 1:500 | Invitrogen |
| Goat anti-rabbit | 568 | 1:500 | Invitrogen |
| DAPI | - | 1:1000 | Sigma |

**Supplementary Table 3.**
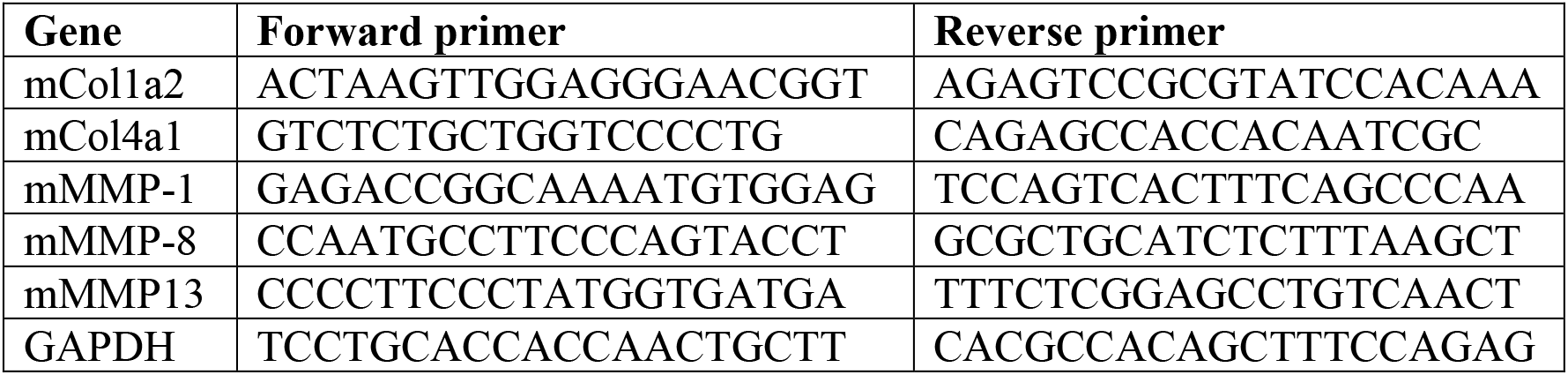
Primer Sequences used for RT-PCR

